# Electroencephalogram recordings in Jhana states: An open dataset

**DOI:** 10.64898/2026.06.27.734949

**Authors:** Marco S Fabus, Stephen Zerfas, Alex Gruver, Maria Fini, Tamaz Gadaev, Kathryn J Devaney

**Affiliations:** School of Medicine and Biomedical Sciences, University of Oxford, Oxford, UK; Jhourney Inc, San Francisco, CA, USA; School of Biological and Health Systems Engineering, Arizona State University, Tempe, AZ, USA; UC Berkeley Center for the Science of Psychedelics, Berkeley, CA, USA

## Abstract

The use of meditation as a tool to improve human wellbeing is receiving considerable scientific interest. However, most existing research has focused on concentration-based practices. One powerful alternative is *jhana* meditation, which leads to states characterised by self-reinforcing bliss, potentially useful for a variety of clinical and scientific domains. However, our understanding of these states is limited by small amounts of data and poor access to experts. To enable new insights, here we release the largest to date and first open-access dataset of electroencephalographic and physiological recordings in expert Jhana meditators. This includes 100+ hours of data in N=26 subjects across three retreats, alongside a detailed description and example code illustrating analysis of the data. This open dataset release can enable wider collaboration and has the potential to move us closer to an understanding of endogenously generated altered states of consciousness.

## Introduction

Meditative practices have received considerable interest for their potential to improve well-being^1^ and provide insights into the workings of the human mind^2^. Existing research has largely focused on concentration-based mindfulness meditation practices, but increasingly a wide variety of practices are being studied^3^. One such practice is *jhana* meditation, a practice that leads to profoundly altered states of consciousness in expert meditators^4^. Briefly, starting from a state of heightened concentration or loving-kindness (approaches vary), this practice, described in Buddhist traditions over 2000 years ago, involves eight sequential jhana stages (J1-J8). The first four jhanas are known as “form jhanas” and the second four (J5-J8) are termed the “formless jhanas” depending on the presence of bodily form sensations. More specifically, focusing on the form jhanas, the first jhana (J1) is characterized by intense physical bliss, often alongside smiling and increased muscle tension. In the second jhana (J2), joy and happiness permeate the body, giving way to a deep contentment in the third jhana (J3) and finally profound equanimity described in the fourth jhana (J4). Such powerfully pleasant states may have important implications for changing a person’s relationship with pleasure, particularly in the context of addiction treatment^5^.

Despite the potential importance of jhana states, neuroscientific understanding of these states remains very limited. One functional magnetic resonance imaging (fMRI) case study of an expert meditator suggested that jhana meditation might represent self-stimulation of the brain’s reward system^6^. A case series of electroencephalogram (EEG) recordings from jhana meditators highlighted visually present spindle/alpha activity as a possible correlate, though this analysis did not test this formally nor split subjects according to self-reported jhana stages^7^. A study of EEG correlates in N=12 meditators in related states of deepening ‘absorption’ observed increased frontal midline theta as well as increased alpha activity^8^, though this study was not explicitly concerned with the jhana tradition. A recent fMRI case study also found distinct brain states associated with jhanas^9^. These results are interesting in light of non-pharmacological altered states of consciousness, which may have promising therapeutic effects with a more favourable side effect profile.

For instance, certain meditation techniques have been shown to acutely increase dopamine tone, as measured by Positron Emission Tomography (PET)^10^.

Overall, the above current jhana evidence is limited to individual case studies and small cohorts. This is likely due to a low number of expert jhana meditators and poor access to them. Whilst the above studies suggest a scientifically fascinating set of states, jhana meditation is difficult to master and expertise can take thousands of hours. To make these states more widely accessible, individualised physiological feedback may aid training. In existing mindfulness-based meditation paradigms, neurofeedback technology is already showing promise in developing such brain training protocols^11–13^.

In this paper, we present, to our knowledge, the largest-ever dataset of brain recordings during jhana states. Specifically, we describe a fully openly available dataset of N=71 EEG and physiological recordings from N=26 expert meditators, collected over three retreats, forming 100+ hours of data from the first four jhanas alongside a variety of baseline and artifact-control recordings. We believe this dataset can lead to a significantly better understanding of neural and physiological activity during the jhanas, and thus in the future inform potential machine learning and neurofeedback applications.

## Materials and Methods

Broadly, this paper presents observational physiological data from N=26 expert meditators (multiple sessions from each participant) across three retreats taking part between March 2023 and June 2023. These are termed Dataset 1 (N=7 subjects), Dataset 2 (N=2 subjects), and Dataset 3 (N=17 subjects). Inclusion criteria for all datasets included being an expert jhana meditator (defined as the ability to reliably access jhana states at will), no history of epilepsy or neurological abnormalities, and no changes to usual medications during the retreats. All subjects had the protocol thoroughly explained, had an opportunity to ask questions, and provided written informed consent before partaking in the study. IRB approval was granted by BRANY IRB for Jhourney Protocol 1337032023. A subject was excluded if they did not reach the jhanas (N=2), or if they subsequently withdrew their consent to participate and share the data (N=1).

### Dataset 1

Observational data was collected in N=7 subjects (age = 41 (26-64), mean (range); 7 males; N=6 right-handed; lifetime meditation hours 4,570 (500-17,700); more details about subjects in the Data Records section below). Data was collected during a jhana-focused retreat in March 2023, hosted by Jhourney, Inc. This retreat taught jhanas in the tradition of Ayya Kemma / Leigh Brasington^14^ with some influence from Thanissaro Bikku / Rob Burbea^15^. In this practice, the breath is the meditation object prior to entering jhana, forming the ‘Access Consciousness’ (AC) stage. The tradition focuses on practicing volitional control over the jhanas, as well as volitional control over the jhanas, as well as the progressive refinement of jhana factors across the four form jhanas—applied thought (*vitakka*), sustained thought (*vic*ā*ra*), rapture (*p*ī*ti*), pleasure (*sukha*), and one-pointedness (*ekaggat*ā)—with each successive jhana characterized by the dropping away of coarser factors as absorption deepens. These traditions treat jhana as a learnable, repeatable skill—entered via pleasure and rapture rather than a visual *nimitta*—and as a stable samatha platform from which insight practice can be conducted.

Before each recording session, participants were instructed to not wear any make-up, hair products, or facial lotion. Then, they completed a pre-session questionnaire, which included questions about their caffeine intake and meditation experience. A 64-channel electroencephalography cap (EEG, BioSemi) was applied with standard international 10-20 electrode positioning. Further electrodes were attached to measure the electrodermal activity (EDA, also known as galvanic skin response, GSR), electrocardiogram (ECG, right collarbone), electrooculogram (EOG, left vertical and right horizontal), and electromyography (EMG, left mastoid / left temple, left zygomaticus, and the laryngeal prominence), as well as a respiration belt (Vernier). The electrodes were chosen to capture potential artifacts, especially smiling which is common in the first jhanas. Data was sampled at 1000Hz with LabRecorder (LSL) and post-hoc down-sampled to 256Hz, with no software filters applied. Where available (detailed in Data Records below), subjects also had pulse oximetry recorded (Wellue SleepU Wrist Pulse Oximeter (Model PO3), Shenzhen Viatom Technology Co., Ltd., Shenzhen, China.). Subjects were also given a computer mouse to indicate transitions between jhana stages with left mouse clicks. Then, the recording session began (mean 5.3 sessions from each subject, range 3-6). Each session was done in two sections.

The aim of the first section was to record several muscle artifacts likely to occur during jhana meditation. For each type of artifact exercise, participants received a verbal instruction and then repeated it three times. Artifact exercises included blinking, looking left then right, rolling eyes up whilst eyes were closed, clenching their jaw, relaxing their tongue, relaxing their face and throat, relaxing their jaw, a big smile, a gentle smile, and swallowing.

The second section was jhana meditation. First, baseline activities were recorded. This included 3 minutes where participants were instructed to plan their day with eyes closed and 3 minutes of a mental arithmetic task with eyes closed (subtracting in intervals of 13 from 1000). Participants were then told to go through access concentration into the jhanas in order. They were instructed to click on arrival in a given stage, click again if the stage gets significantly deeper, and double click if something abnormal happens. After 20 minutes in access concentration, participants were told to move towards Jhana 1. Once they indicated arrival by clicking, 10 minutes were timed and participants were told to move towards Jhana 2. This was repeated with Jhana 3 and Jhana 4. If participants did not reach Jhana 4 within 5 minutes of the instruction, or after Jhana 4, they were told to relax and let the experimenter know when they were ready to record the post-meditation baseline exercises. Once ready, similarly to the baseline exercises, 3-minute eyes-closed recordings were made of day planning and an arithmetic exercise.

### Dataset 2

Here, observational data was collected in N=2 subjects (age = 27 and 49; 1 female / 1 male, both right-handed; lifetime meditation hours 1,800 and 25,000). Data was collected during a jhana-focused retreat in November 2022. This retreat was taught by a student of Leigh Brasington. The methodology was similar to that of Dataset 1.

### Dataset 3

Final observational data was collected in N=17 subjects (age = 56 (26-78); 12 males / 5 females; all right-handed; lifetime meditation hours 9,740 (300-67,000)). Data was collected during a jhana-focused retreat in May 2023, organised as part of the “Dependent Origination Symposium”. This retreat taught jhanas in the tradition of “Tranquil Wisdom Insight Meditation” (TWIM). Broadly, TWIM uses loving-kindness (*metta*^16^) instead of the breath as the meditation object before jhanas (no access concentration step), and less focus is put on volitional control of jhanas, focusing instead on the natural movements of the nervous system during a meditation sitting.

Again, broadly, the methodology was identical to Dataset 1, with minor differences. The first difference was that a game controller was used instead of a mouse to indicate jhana transitions. In the artifact exercise section, eyebrow flexing was added and smiling was instructed to be held for 5 seconds. Pre-sit baselines included eyes-closed mind wandering, and eyes-closed mind wandering whilst smiling. During the jhana section, participants were free to decide how much time to spend in each jhana, but the overall sitting was timed to 60 minutes, after which subjects were told to relax and move to the post-meditation baselines. Participants indicated arrival in a given jhana by pressing *A* on the game controller *K* times for the *K*th jhana, and there was no access concentration stage. Post-sit baselines were the same as pre-sit baselines.

For dissemination, all data was standardized into the BIDS-EEG format. EEG was re-referenced to the average reference.

### Example EEG analysis

We undertook a proof-of-concept example analysis of the EEG data to demonstrate its potential and illustrate its use to other researchers. Specifically, alpha power spectral density (8-13Hz) was extracted for each electrode in each subject and each experimental stage using Welch’s method as implemented in the MNE psd_array_welch function. The median topographical map across subjects was calculated. A jupyter notebook with all code required to replicate this analysis is provided alongside the dataset. This notebook demonstrates how to load the data, extract relevant experimental stages, and apply EEG analysis methods.

## Data Records

The dataset contains N=71 session from N=26 subjects in total. The dataset is organised in line with the Brain Imaging Data Structure (BIDS) specifications for EEG^17^. Each subject has its own folder, with each session as a child folder to the subject. Each session folder includes the physiological data in the European Brain Data Format (.edf). As well as this, there are .tsv files with channel and event descriptions and a JSON file describing further details of the recording. Example code to use the data is provided in a Jupyter notebook. More details of all datasets can be found in Tables 1-4 below. A full description of all datasets, baselines, minutes, per jhana, and notes on individual recordings can be found in the dataset download folder (see Code availability below).

**Table 1:** Summary of the datasets.

| Dataset | N (people) | N (EEG sessions) | N (EEG hours) |
| --- | --- | --- | --- |
| Dataset 1 | 7 | 37 | 57.6 |
| Dataset 2 | 2 | 2 | 2.66 |
| Dataset 3 | 17 | 32 | 48.4 |

**Table 2:** Overview of Dataset 1.

| TWIM Dataset | N (people) | Demographics |  | Dataset |  |  |  |
| --- | --- | --- | --- | --- | --- | --- | --- |
|  |  | Age | Sex | Sessions per subject | Total sessions | Total hours | Mins per jhana |
| EEG | 7 | 41 [26-64] | 7M | 5 [3-6] | 37 | 57.6 | 249 |
| EOG/ECG/EMG | 7 | 41 [26-64] | 7M | 5 [3-6] | 37 | 57.6 | 249 |
| EDA (GSR) | 7 | 41 [26-64] | 7M | 5 [3-6] | 37 | 57.6 | 249 |

**Table 3:** Overview of Dataset 2.

| TWIM Dataset | N (people) | Demographics |  | Dataset |  |  |  |
| --- | --- | --- | --- | --- | --- | --- | --- |
|  |  | Age | Sex | Sessions per subject | Total sessions | Total hours | Mins per jhana |
| EEG | 2 | 27, 49 | 1F, 1M | 1 | 2 | 2.66 | 9.3 |
| EOG/ECG/EMG | 2 | 27, 49 | 1F, 1M | 1 | 2 | 2.66 | 9.3 |
| EDA (GSR) | 2 | 27, 49 | 1F, 1M | 1 | 2 | 2.66 | 9.3 |

**Table 4:** Overview of Dataset 3.

| TWIM Dataset | N (people) | Demographics |  | Dataset |  |  |  |
| --- | --- | --- | --- | --- | --- | --- | --- |
|  |  | Age | Sex | Sessions per subject | Total sessions | Total hours | Mins per jhana |
| EEG | 17 | 56 [26-78] | 5F, 12M | 2 [1-3] | 32 | 48.4 | 303 |
| EOG/ ECG/ EMG | 17 | 56 [26-78] | 5F, 12M | 2 [1-3] | 32 | 48.4 | 303 |
| EDA (GSR) | 11 | 56 [26-78] | 5F, 12M | 1 [1-2] | 14 | 21.4 | 138 |

We also carried out a proof-of-concept EEG analysis of this data. Specifically, we extracted alpha power across each experimental stage (Figure 1). The average across all stages and subjects showed a typical EEG topography with dominant occipital alpha oscillations. Interestingly, different jhana stages appeared to have distinct alpha topographies, though this was not formally tested.

**Figure 1:**
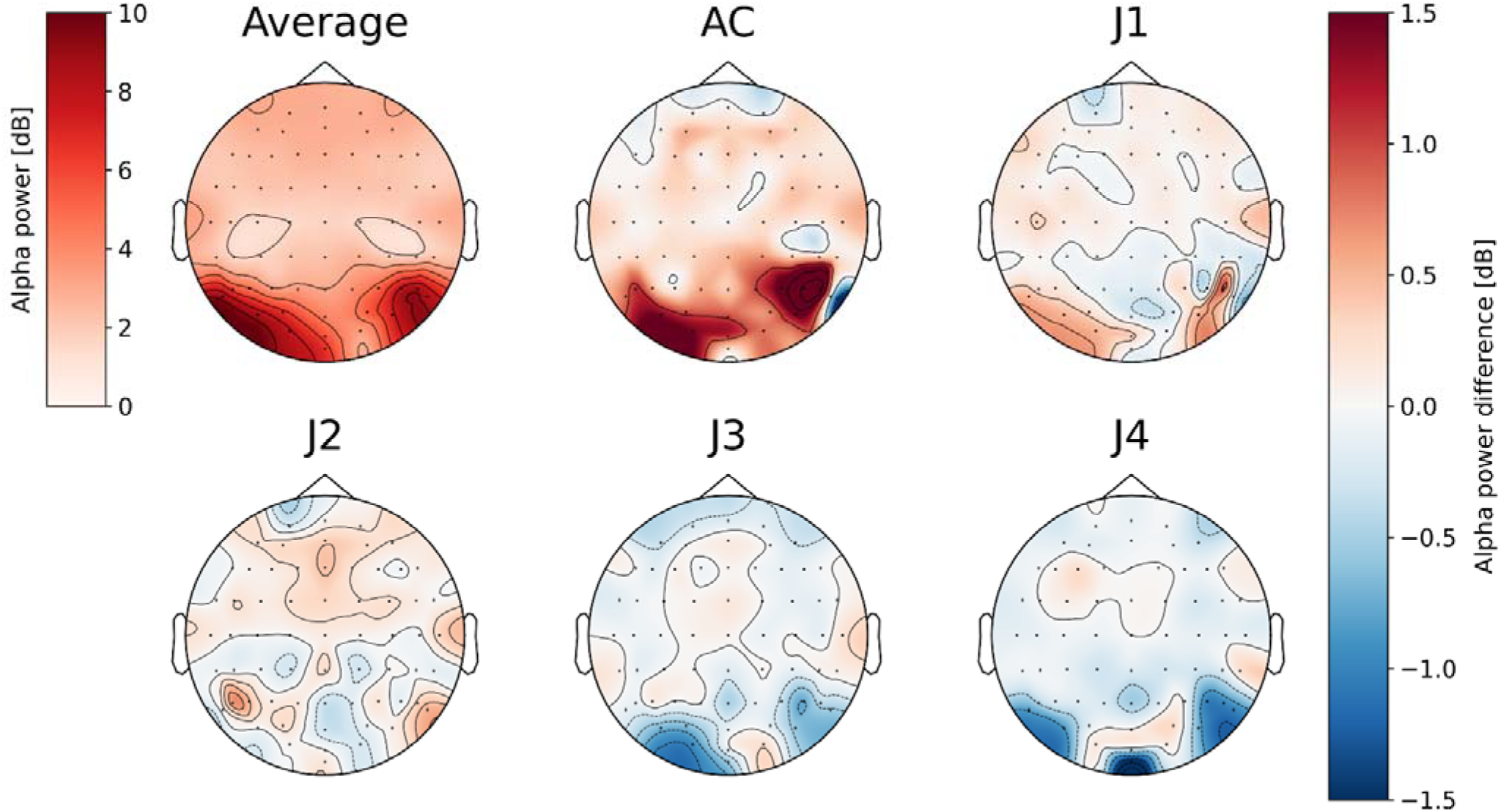
Alpha power topography across the experiment (median across subjects). Top left – median across the whole experiment. Rest of the panels show demeaned alpha power in relevant jhana stages.

## Code availability

A full repository with instructions on how to access the data as well as example scripts can be found at https://github.com/tamazgadaev/jhana_eeg. The specific address for direct data download is https://drive.google.com/drive/folders/1PM5lDLJr2LKleADL9zM65HOQY8ykYg2_?us_p=drive_link. For specific queries, please contact.

## Acknowledgements

Authors would like to extend thanks to Tucker Peck, Leigh Braisington, Delson Armstrong, and Michael Taft for help with participant recruitment.

MSF was supported by the Wellcome Centre for Integrative Neuroimaging Wellcome Trust Studentship [Grant number 203139/Z/16/Z], KJD is supported by the Ad Astra Chandaria Foundation. For the purpose of open access, the authors have applied a CC-BY public copyright license to any Author Accepted Manuscript version arising from this submission.

